# Host brain infection induces RNAi epimutation in *Mucor circinelloides* and enhances blood-brain barrier invasion

**DOI:** 10.64898/2026.10.23.761329

**Authors:** María Isabel Navarro-Mendoza, Carlos Pérez-Arques, Yeseul Choi, Emma Cao, Zanetta Chang, Ziyan Xu, Yong-Sun Bahn, E. Ashley Moseman, Joseph Heitman

## Abstract

Mucorales fungi cause mucormycosis, a rapidly progressive invasive fungal disease with high mortality, particularly in immunocompromised patients and in severe disseminated infections. Central nervous system involvement is among the most devastating manifestations of mucormycosis, yet the mechanisms that enable Mucorales to adapt to the brain environment remain poorly understood. In *Mucor* species, RNA interference (RNAi) can generate reversible heritable epimutations that silence specific genes without altering DNA sequence, providing a transient and reversible mechanism for antifungal drug resistance. Here, we show that host brain infection induces RNAi-dependent silencing of endogenous fungal loci that promote central nervous system adaptation. Small RNA sequencing of *Mucor circinelloides* recovered from infected mouse organs identified seven brain epimutation loci, named *bepA* to *bepG*, that accumulated small RNAs preferentially after brain infection. Among these, *bepA* encodes a conserved mitogen-activated protein kinase (MAPK)-like serine/threonine kinase related to cell wall integrity signaling. Deletion of the *bepA* gene confers resistance to the calcineurin inhibitors FK506 and cyclosporin A, allowing hyphal growth under conditions that impair calcineurin-dependent morphogenesis. BepA modulates FK506-induced MAPK signaling and controls a transcriptional program enriched in cell-wall remodeling genes. To model fungal central nervous system entry by a filamentous fungal pathogen, we established a human blood-brain barrier transwell platform that quantifies *Mucor* migration across brain endothelial monolayers. This model revealed that *bepA* mutation enhances fungal migration across brain endothelial monolayers without detectable changes in electrical barrier resistance. These findings identify brain-induced RNAi epimutation as a mechanism of organ-specific fungal adaptation and reveal reversible silencing of a MAPK-like kinase as a route to enhanced central nervous system invasion.

## Introduction

Mucormycosis is a rapidly progressive invasive fungal disease characterized by angioinvasion, tissue necrosis, and mortality rates that can exceed 50 to 90 % in disseminated cases. The infection predominantly affects immunocompromised patients, especially individuals with diabetes, hematologic malignancies, transplantation, or corticosteroid-induced immunosuppression. The treatment armamentarium is strikingly limited, relying mainly on surgical debridement and aggressive antifungal therapy with only three effective drugs: amphotericin B, posaconazole, and isavuconazole (1). The World Health Organization recently designated the Mucorales among the high-priority fungal pathogens, underscoring the urgent need to understand mechanisms governing their virulence and antifungal resistance mechanisms (2).

One of the most severe clinical manifestations of mucormycosis is rhino-orbital-cerebral disease, which arises following inhalation of spores into the paranasal sinuses and subsequent contiguous invasion of the orbit, eye, and brain (3). Central nervous system (CNS) involvement is common in both this form and with disseminated disease, and is associated with a particularly high mortality rate (4). These clinical manifestations reflect the marked angioinvasive capacity of the Mucorales, enabling vascular thrombosis, tissue necrosis, and hematogenous or direct extension into the brain. Although molecular mechanisms mediating endothelial invasion and angioinvasion have been identified, these pathways do not explain how the Mucorales adapt to the unique microenvironment of the CNS.

The Mucorales exhibit intrinsic resistance to many antifungal agents, including short-tailed azoles and the echinocandins. Beyond stable genetic mechanisms, these fungi possess an RNA interference (RNAi) pathway that enables reversible gene silencing in response to stress. This RNAi-based epimutation was first described as a mechanism of antifungal drug resistance in *Mucor* species, where exposure to drugs such as FK506 or 5-fluoroorotic acid induces the production of locus-specific small RNAs that direct degradation of target transcripts, and silence drug target genes without altering their DNA sequence (5, 6). This silencing confers transient resistance that can revert upon removal of selective pressure, providing a flexible, non-Mendelian mode of inheritance mediated by small RNAs (7).

In addition to drug resistance, the RNAi machinery in the Mucorales contributes to aspects of genome regulation and pathogenesis. RNAi suppresses transposable elements and contributes to genome stability and centromere identity (8). Disruption of RNAi pathways alters virulence traits and impairs adaptation during host immune interaction, indicating that RNAi contributes to fungal fitness *in vivo* (9). Importantly, the frequency and stability of RNAi-dependent epimutations are influenced by the host environment. Experimental infection studies demonstrated that epimutation frequency increases following *in vivo* passage compared to *in vitro* growth, indicating that host-associated conditions stimulate RNAi silencing. Isolates recovered from the brain exhibit higher rates of epimutation and prolonged persistence compared to isolates from other organs. In these models, brain passage enhanced the emergence of epimutations conferring antifungal drug resistance (10). These findings suggest that host niches can modulate RNAi activity and thereby influence fungal gene expression programs during infection.

To date, RNAi-dependent epimutations in the Mucorales have been characterized primarily as mechanisms targeting antifungal drug resistance genes. Whether infection-induced RNAi activity extends to regulatory signaling pathways that control host-stress adaptation and tissue invasion has not been determined. Given the organ-specific dynamics of epimutation following *in vivo* passage, host niches could bias silencing toward genes that control host interaction.

Here, we identify brain-associated small RNA accumulation at a mitogen-activated protein kinase (MAPK)-like gene in *M. circinelloides*. Loss of this kinase rewires calcineurin signaling, alters cell wall remodeling programs, and enhances migration across a human blood-brain barrier (BBB) model. These findings establish host-induced RNAi epimutations as a regulatory mechanism of fungal adaptation during central nervous system infection.

## Results

### Brain passage induces RNAi-based silencing at multiple epimutation loci

To determine whether distinct host tissue environments promote RNAi-based epigenetic adaptation during infection, we performed small RNA sequencing of *Mucor circinelloides* recovered from infected organs. Mice were infected intravenously with *M. circinelloides* 1006PhL, a human isolate. After a 48-hour infection period, fungal colonies were isolated from brain and liver, followed by plating and outgrowth under standardized conditions for 4, 6, and 8 days prior to RNA purification and small RNA profiling (Fig. 1A).

**Figure 1.**
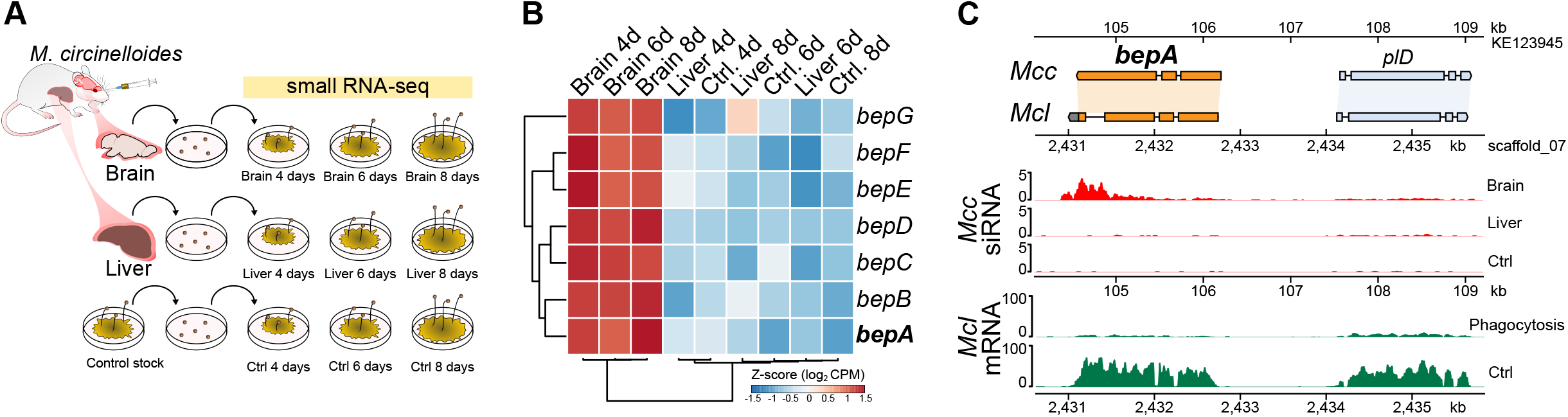
Brain passage induces RNAi-based epimutation of *bepA*. **(A)** Experimental design for small RNA sequencing (small RNA-seq). Mice were infected intravenously with *Mucor circinelloides* spores, and brain and liver tissues were recovered. For each organ and time point, recovered fungal material was plated, grown for 4, 6, or 8 days, and used for RNA extraction and small RNA sequencing. An in vitro control stock was processed in parallel. **(B)** Heatmap depicting brain epimutation (*bep*) genes (*bepA* to *bepG*) that show increased accumulation of small RNAs in the indicated conditions. Values are shown as Z-score normalized log2 counts per million (CPM) derived from small RNA-seq, and hierarchical clustering was applied to visualize similarities across samples. Increased small RNA abundance is interpreted as increased RNAi-associated silencing of the corresponding locus in that condition. **(C)** Conserved synteny at the *bepA* locus in *M. circinelloides* (Mcc) and *M. lusitanicus* (Mcl), and infection-associated expression patterns across the locus. Upper panels show the genomic organization of *bepA* and the adjacent phospholipase D (*pld*) gene in both species. Red panels show small RNA accumulation across the *bepA* region in *M. circinelloides* recovered from brain, liver, and control conditions. Green panels show mRNA expression across the syntenic region in *M. lusitanicus* during interaction with mouse macrophages compared to control conditions.

Genome-wide analysis of small RNA accumulation revealed that brain passage induces a specific RNAi response in *M. circinelloides*. We identified a discrete set of seven loci that accumulate small RNAs following brain infection in mice (named *bep*, for <u>b</u>rain <u>ep</u>imutation, Table 1). These loci include one encoding a MAPK-like kinase, *bepA*, and six uncharacterized genes (designated *bepB-bepG*). *bep* loci exhibited increased small RNA abundance specifically in brain-derived samples, consistent with enhanced RNAi-mediated silencing in this host niche (Fig. 1B). This enrichment was observed across all analyzed post-recovery culture time points. In contrast, liver-derived and *in vitro* control samples displayed minimal small RNA signals at these loci.

**Table 1.** *bep* genes and predicted protein domains.

| Gene | ID | Predicted domains |
| --- | --- | --- |
| *bepA* | HMPREF1544_04416 | PF00069 Protein Kinase |
| *bepB* | HMPREF1544_00249 | pentatricopeptide repeats involved in protein-RNA interactions |
| *bepC* | HMPREF1544_08566 | pentatricopeptide repeats involved in protein-RNA interactions |
| *bepD* | HMPREF1544_04950 | WW protein-protein interaction domain |
| *bepE* | HMPREF1544_11569 | Unknown |
| *bepF* | HMPREF1544_09922 | Unknown |
| *bepG* | HMPREF1544_11570 | Unknown |

Among these loci, *bepA* showed robust and consistent small RNA accumulation in brain-derived samples. Unlike the other *bep* loci, which were associated with repetitive genomic regions, *bepA* mapped to a discrete protein-coding locus and encodes a protein with a conserved kinase domain, suggesting a regulatory function and prompting its further characterization as a candidate regulator linking host niche signals to adaptive transcriptional control during infection. Comparative synteny analysis revealed that *bepA* is conserved in *M. circinelloides* and *M. lusitanicus*, maintaining synteny with adjacent genes, including a neighboring phospholipase D (*plD*) locus (Fig. 1C). Mapping of small RNAs across this region in *M. circinelloides* demonstrated infection-associated accumulation limited to the *bepA* locus in brain-derived samples, whereas liver-derived and *in vitro* control samples showed substantially reduced signal. This small RNA accumulation was localized rather than globally distributed, supporting locus-specific regulation rather than repeat-derived RNAi activation.

To further explore whether this locus responds to host interaction in an independent system, we examined transcriptional profiles across the syntenic region in *M. lusitanicus* during interaction with murine macrophages (11) (Fig. 1C). This analysis revealed coordinated changes in *bepA* transcript abundance, showing reduced messenger RNA levels in response to phagocytosis. Together, these observations suggest that repression of the *bepA* gene may contribute to fungal adaptation during infection and could promote virulence in specific host niches. These findings support a model in which host microenvironments selectively activate RNAi silencing of regulatory loci, including *bepA* during infection.

### BepA is a MAPK-like kinase required for calcineurin-dependent signaling

To define the functional role of BepA, we generated *bepA* deletion (*bepA*Δ) and overexpression (*bepAOE*) strains (Fig. S1). We then assessed growth of these strains under a panel of antifungal stresses relevant to mucormycosis treatment and signaling, including calcineurin inhibitors (FK506 and cyclosporin A), the TOR inhibitor rapamycin, ergosterol-targeting drugs (amphotericin B and posaconazole), and the cell wall inhibitor caspofungin (Fig. S2). In the Mucorales, calcineurin activity is required for hyphal growth, and inhibition of calcineurin results in a stable yeast morphology under aerobic conditions (12). This morphogenetic transition is antagonistically linked to activation of the cAMP-PKA pathway, whereas anaerobic yeast growth is driven primarily through PKA activation independent of calcineurin function (13). Sequence analysis revealed that *bepA* encodes a predicted MAPK containing the conserved glycine-rich loop, HRD catalytic motif, DFG motif, and a canonical TEY activation loop sequence, consistent with a cell wall integrity MAPK of the Mpk1/Slt2 family. Among the conditions tested, independent *bepA*Δ mutant strains exhibited increased resistance to calcineurin inhibitors, maintaining hyphal growth, whereas sensitivity to amphotericin B, posaconazole, and caspofungin was comparable to wild type (Fig. 2A and Fig. S2A). In contrast, *bepA*Δ mutant strains displayed normal yeast growth under anaerobic conditions (Fig. S2B), indicating that BepA is dispensable for PKA-driven yeast growth but required for full calcineurin-dependent dimorphic responses. These results support a specific role of BepA in calcineurin-dependent signaling.

**Figure 2.**
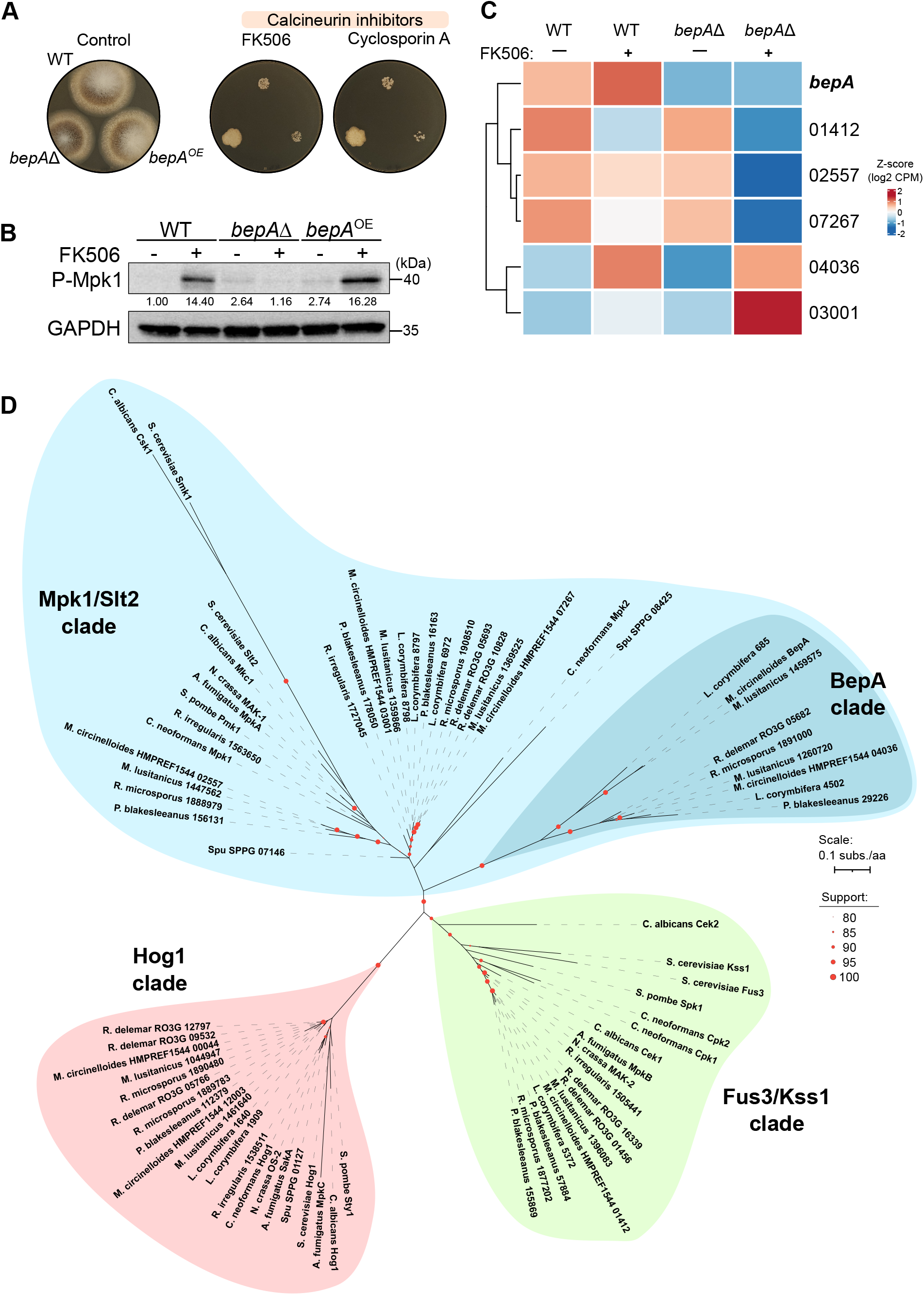
BepA is a conserved MAPK-like kinase required for calcineurin-dependent signaling. **(A)** Growth of wild-type (WT), *bepA* deletion (*bepA*Δ), and *bepA* overexpression (*bepA*^OE^) strains on YPD control medium and in the presence of the calcineurin inhibitors FK506 and cyclosporin A. The *bepA*Δ strain displays increased resistance to FK506 compared to WT. **(B)** Western blot analysis of MAPK phosphorylation in WT, *bepA*Δ, and *bepA*^OE^ strains grown in the absence or presence of FK506. A phospho-MAPK signal (labeled P-Mpk1 in the panel) was detected with an anti-phospho-p44/42 MAPK antibody, and GAPDH served as a loading control. Band intensities were quantified with ImageJ, normalized to GAPDH, and expressed relative to WT without FK506. FK506 induces MAPK phosphorylation in the WT strain, whereas this response is strongly reduced in the *bepA*Δ mutant. **(C)** Heatmap showing differential expression of *bepA* and selected cell wall integrity-associated genes in WT and *bepA*Δ strains grown in the absence or presence of FK506. Expression values are shown as row-scaled Z-scores of log2 CPM values. **(D)** Maximum-likelihood phylogeny of conserved protein kinase domains from BepA and MAPK candidates identified across representative Mucorales, together with homologs from *Rhizophagus irregularis*, *Spizellomyces punctatus*, and characterized fungal MAPKs representing the Mpk1/Slt2 cell wall integrity, Hog1 stress-response, and Fus3/Kss1 pheromone/filamentation pathways. Node support values are shown as SH-aLRT/ultrafast bootstrap percentages based on 1,000 replicates each; branch lengths represent amino-acid substitutions per site. The tree is shown unrooted.

In the fungal human pathogen *Cryptococcus neoformans,* calcineurin inhibition results in increased phosphorylation of the cell wall integrity MAPK Mpk1, leading to the activation of the cell wall integrity response (14). To determine whether calcineurin inhibition similarly induces MAPK phosphorylation in *M. circinelloides,* and whether BepA contributes to this response, we examined Mpk1 phosphorylation following FK506 treatment. We employed an anti-phospho-p44/42 MAPK antibody (phospho-Mpk1) to assess phosphorylation in the wild-type, *bepA*Δ mutant, and *bepA*^OE^ strains. Similar to *C. neoformans*, Mpk1-like phosphorylation was barely detectable under basal conditions but was strongly induced by FK506 in the wild-type strain. In contrast, *bepA*Δ mutants failed to show FK506-induced Mpk1 phosphorylation, whereas the *bepA^OE^* strain retained a strong phospho-Mpk1 signal (Fig. 2B, Fig. S3). Densitometric analysis of relative phospho-Mpk1 MAPK levels indicated that BepA is required for full Mpk1 phosphorylation in response to calcineurin inhibition. Consistent with this signaling role, transcriptomic analysis revealed that FK506 treatment induced *bepA* expression in the wild-type strain, whereas this response was absent in the *bepA*Δ mutants (Fig. 2C). These data position BepA as a kinase required for efficient engagement of the cell wall integrity MAPK pathway during calcineurin inhibition.

Because BepA contains a conserved TEY activation motif, the phospho-Mpk1 signal can be interpreted in two non-mutually exclusive ways. In one model, BepA may act upstream of, or in parallel with, the canonical cell wall integrity MAPK module and is required for FK506-induced phosphorylation of Mpk1. In an alternative model, BepA itself may correspond to the phospho-Mpk1 species detected by this antibody or may represent the principal Mpk1 activated upon calcineurin inhibition. The strongly reduced signal in the *bepA*Δ mutant strain and retained signal in the *bepA*^OE^ strain are consistent with both possibilities. To examine the evolutionary relationship of BepA to canonical fungal MAPKs, we performed a broad phylogenetic analysis of MAPK kinase domains across Mucorales and representative fungal lineages (Fig. 2D). Candidate kinases were identified based on sequence similarity and MAPK-family profile searches, filtered for the presence of a conserved protein kinase domain, and analyzed together with experimentally characterized representatives of the three major fungal MAPK families, including the Mpk1/Slt2 cell wall integrity (CWI), Hog1 stress-response, and Fus3/Kss1 pheromone/filamentation pathways.

This phylogenetic analysis argues against BepA being the canonical Mucorales Mpk1 ortholog. A distinct set of Mucorales proteins clustered with established Mpk1/Slt2 cell wall integrity MAPKs, including a CWI lineage that might represent the Mucorales canonical Mpk1 orthologs and a separate CWI clade closely related to *C. neoformans* Mpk2. In contrast, BepA grouped within a strongly supported MAPK lineage restricted to Mucorales in our taxon sampling, with closely related homologs in *Mucor*, *Rhizopus*, *Lichtheimia*, and *Phycomyces* species. No clear representative of this clade was detected in the sampled Glomeromycota, Chytridiomycota, or Dikarya. Together, these results support BepA as an Mpk1-related, functionally specialized paralog that emerged during Mucorales evolution rather than as the direct ortholog of canonical Mpk1.

Together, the phylogeny and functional data support a model in which BepA is not the canonical Mpk1 ortholog, but a Mucorales-specific MAPK paralog that arose through lineage-specific expansion of an ancestral Mpk1-related family and subsequently acquired a specialized role in the CWI stress signaling pathway and host adaptation.

### BepA regulates cell wall remodeling genes

To define the transcriptional programs controlled by BepA activity, we performed RNA sequencing in wild-type, *bepA*Δ mutant, and *bepA^OE^* strains in the presence or absence of the calcineurin inhibitor FK506. Differential expression analysis revealed extensive transcriptional remodeling in response to calcineurin inhibition in the wild-type strain (Fig. 3A). These FK506-responsive genes reflect the broad transcriptional changes associated with calcineurin inhibition in *Mucor*. In parallel, comparison of *bepA^OE^* and *bepA*Δ mutant strains identified a distinct set of BepA-dependent genes (Fig. 3B). A large fraction of BepA-dependent genes (607 of 775) overlapped with those responsive to FK506 treatment, indicating that BepA controls a subset of calcineurin-regulated transcriptional outputs (Fig. 3C). This overlap supports a model in which BepA acts as a key signaling node integrating calcineurin inhibition into downstream transcriptional responses.

**Figure 3.**
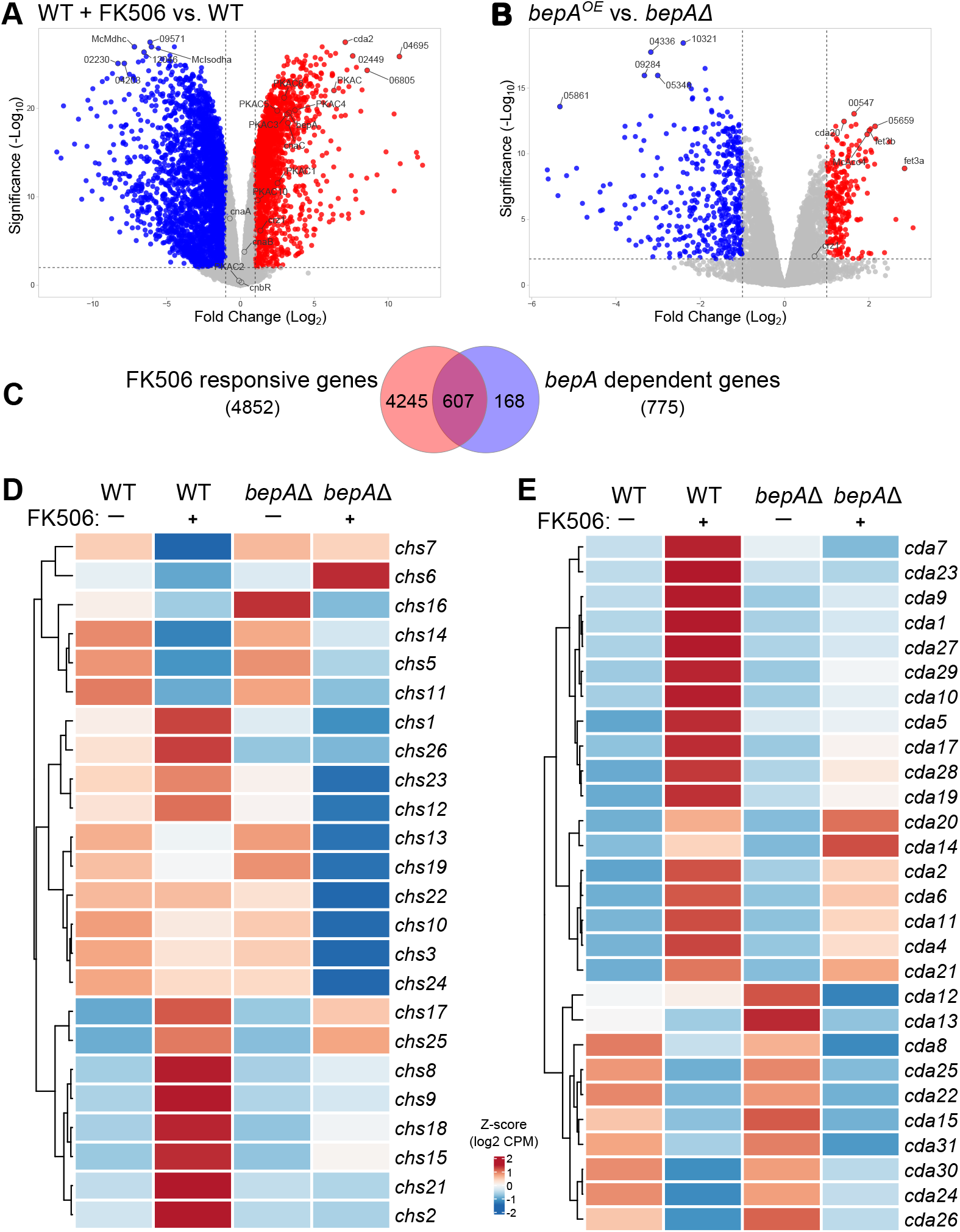
BepA-dependent transcriptional responses to calcineurin inhibition include cell wall remodeling genes. **(A)** Volcano plot showing differentially expressed genes in the wild-type (WT) strain grown in the presence of FK506 compared to untreated WT cells. Significantly upregulated and downregulated genes are indicated in red and blue, respectively. **(B)** Volcano plot showing differentially expressed genes between *bepA*^OE^ and *bepA*Δ strains. Genes exhibiting significant changes in expression are highlighted in red and blue. **(C)** Venn diagram showing the overlap between FK506-responsive genes and *bepA*-dependent genes. **(D)** Heatmap showing expression profiles of chitin synthase genes (*chs*) in WT and *bepA*Δ strains grown in the absence or presence of FK506. **(E)** Heatmap showing expression profiles of chitin deacetylase genes (*cda*) in WT and *bepA*Δ strains grown in the absence or presence of FK506. In both heatmaps, values represent Z-scores of log_2_ transformed counts per million (CPM).

Gene ontology (GO) and functional enrichment analyses of the overlapping FK506- and BepA-dependent gene sets revealed a pronounced enrichment for cell wall biosynthesis and remodeling pathways. Genes involved in chitin metabolism and cell wall architecture were significantly enriched. The Mucorales cell wall is enriched in chitin and more importantly, its deacetylated form chitosan, with relatively low β-glucan content compared to other pathogenic fungi (15). Regulation of chitin synthesis and deacetylation is therefore central to cell wall remodeling during stress and host interaction (16). Expression profiling of chitin synthase (*chs*) genes demonstrated coordinated, BepA-dependent regulation under FK506 treatment (Fig. 3D). Multiple *chs* genes that responded to FK506 in wild-type cells failed to be similarly induced or repressed in the *bepA*Δ mutant background. Similarly, chitin deacetylase (*cda*) genes exhibited altered expression patterns in *bepA*Δ mutant strains (Fig. 3E), further supporting a role for BepA in controlling cell wall architecture during stress. Together, these data demonstrate that BepA integrates calcineurin signaling with downstream transcriptional programs governing cell wall remodeling. These findings further suggest that fine-tuned regulation of BepA contributes to the coordinated expression of multiple *chs* and *cda* paralogs, allowing adjustments in cell wall remodeling in response to environmental stress. In the context of brain infection, RNAi-mediated silencing of bepA could shift the balance of these regulatory programs, potentially contributing to fungal adaptation to the host environment. Given the importance of cell wall architecture in host interaction, we next examined whether loss of *bepA* affects fungal pathogenesis.

### Loss of *bepA* enhances fungal migration across a human blood-brain barrier model

Given that *bepA* silencing was enriched in brain-infected samples and that BepA controls cell wall remodeling programs, we next asked whether loss of BepA alters fungal CNS invasion. To model CNS entry *in vitro*, we established an *in vitro* blood-brain barrier (BBB) model for mucormycosis, adapting a previously validated human BBB transwell system developed for yeast pathogen migration (17, 18). Human brain microvascular endothelial cells (hCMEC/D3) were cultured to confluence on collagen-coated transwell inserts to generate a polarized endothelial monolayer (Fig. 4A). Fungal spores were added to the apical (vascular) compartment, and their migration to the basolateral (brain-proximal) chamber was quantified by plating and counting fungal colonies. To assess whether fungal infection compromises endothelial barrier integrity, transendothelial electrical resistance (TEER) was measured before and after infection. We first established and optimized migration conditions with the wild-type strain by adjusting spore inoculum and incubation time for migration without TEER change, and we then quantified fungal migration of the wild-type strain compared to the *bepA*Δ mutant and *bepA^OE^* strains (Fig. 4B).

**Figure 4.**
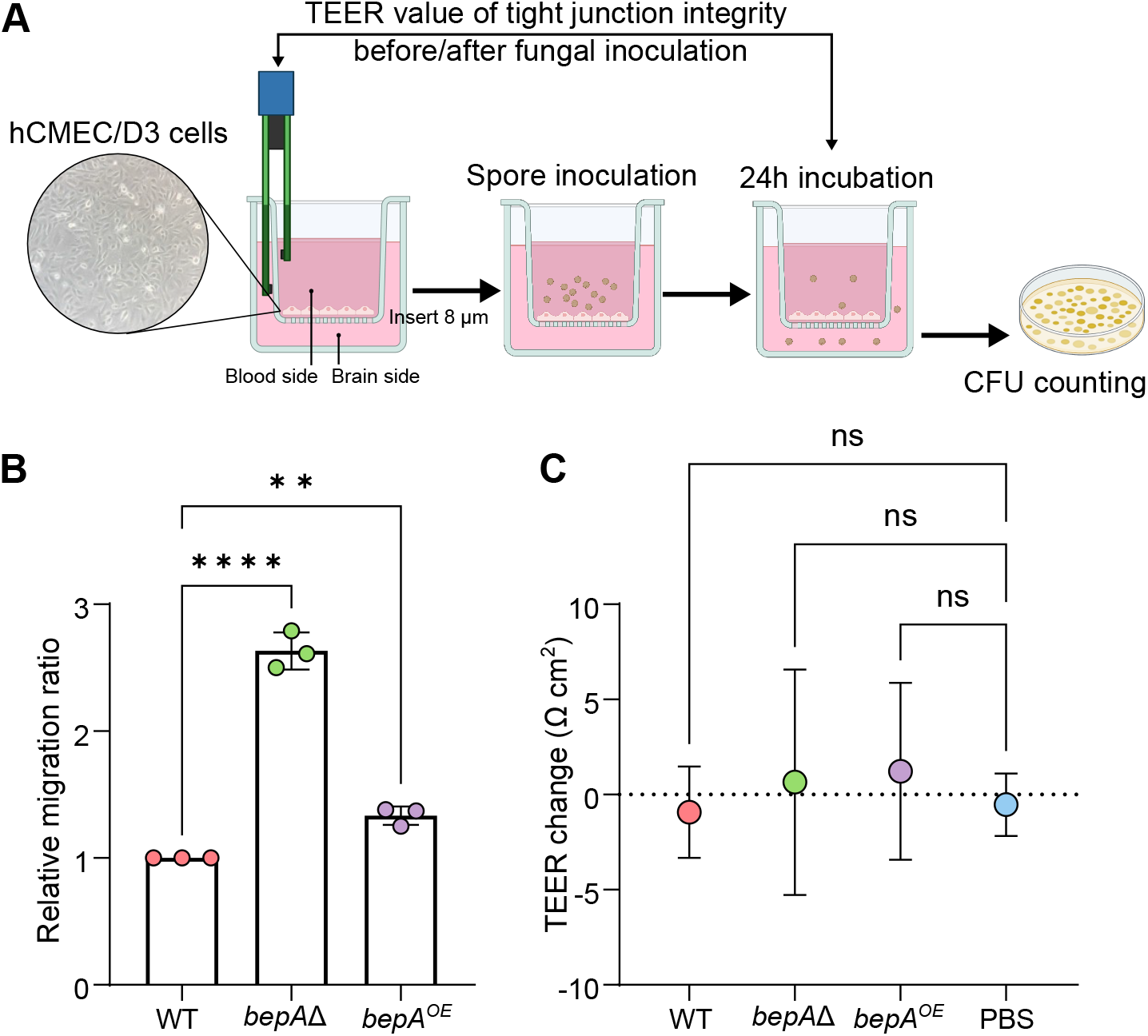
Loss of *bepA* enhances migration across an in vitro blood-brain barrier model. **(A)** Schematic of the in vitro blood-brain barrier (BBB) model using hCMEC/D3 human brain microvascular endothelial cells cultured on transwell inserts. Transendothelial electrical resistance (TEER) was measured before and after fungal inoculation to assess electrical barrier resistance. Spores were added to the upper chamber (blood side), and migration to the lower chamber (brain side) was quantified by collecting and plating fungal biomass, and counting CFU after 24 h. The schematic illustration was created using BioRender. **(B)** Relative migration rate of the wild-type (WT), *bepA*Δ, and *bepA*^OE^ strains across the BBB model. Each dot represents an independent biological replicate (n = 3 per group), and bars indicate mean ± SD. Statistical significance was assessed by one-way ANOVA followed by Tukey’s multiple-comparison test for the indicated comparisons. Statistical significance is indicated (**P < 0.01; ****P < 0.0001). **(C)** Changes in TEER values following fungal inoculation with WT, *bepA*Δ, and *bepA*^OE^ strains, or PBS control (n = 3 per group). Error bars indicate SD. Statistical significance was assessed by one-way ANOVA followed by Tukey’s multiple-comparison test fo the indicated comparisons; ns, not significant.

Deletion of *bepA* significantly increased fungal migration across the endothelial barrier by approximately 2.6- and 2-fold compared to wild-type and *bepA^OE^* strains, respectively (Fig. 4B). TEER measurements before and after infection revealed no significant changes in endothelial barrier resistance (Fig. 4C), suggesting that increased migration was not associated with detectable barrier disruption. These data indicate that loss of BepA enhances transendothelial migration without compromising barrier integrity. This phenotype provides a functional explanation to the enrichment of *bepA* epimutation observed in mouse brain, suggesting that *bepA* gene silencing may confer a selective advantage during CNS invasion.

To determine whether BepA influences virulence *in vivo*, we first assessed survival in a disseminated infection model with cyclophosphamide-immunosuppressed BALB/c mice infected intravenously with wild-type, *bepA*Δ, or *bepA^OE^* strains. Survival curves did not differ significantly between strains (Fig. S4), indicating that loss or overexpression of BepA does not measurably alter overall mortality in this mouse model. To further examine potential effects on fungal dissemination and host responses, we quantified fungal burden and immune cell populations in infected C57BL/6 mice (Fig. S5). Mice infected with the *bepA*Δ mutant strain displayed significant increases in fungal burden in the spleen and brain compared with wild-type controls (Fig. S5). Consistent with this fungal burden, flow cytometric analysis of splenic immune populations revealed increased numbers of neutrophils and inflammatory monocytes in mice infected with the *bepA*Δ mutant (Fig. S5). Together, these results indicate that while BepA does not significantly affect host survival in the BALB/c infection models, its loss alters fungal dissemination and is associated with enhanced splenic immune recognition during infection.

## Discussion

Together, our findings support a model in which the host brain actively drives epigenetic adaptation during fungal infection. RNAi-dependent epimutation in Mucorales has primarily been viewed as a mechanism that generates transient antifungal drug resistance through reversible silencing of drug target genes (5, 6, 19). Our results expand this paradigm by demonstrating that host infection itself can induce RNAi-dependent silencing of endogenous genes that promote tissue-specific adaptation. Rather than merely selecting pre-existing epigenetic variants, the brain emerges as a specialized host niche that promotes a defined RNAi response directed toward endogenous regulatory loci.

This model is consistent with earlier work showing that RNAi is not restricted to genome defense or drug resistance in the Mucorales but also regulates endogenous transcripts that influence fungal development and host interaction. *M. circinelloides* produces exon-derived endogenous small RNAs that regulate protein-coding genes (20), while disruption of the canonical RNAi machinery broadly alters gene expression and affects developmental processes including sporulation, autolysis, sexual interaction, and adaptation to environmental stress (21). In parallel, the non-canonical RNAi pathway controls extensive transcriptional networks required for macrophage adaptation, virulence, and genome stability (9). Together, these studies established RNAi as a versatile mechanism for endogenous gene regulation and environmental adaptation. Our findings extend this framework by revealing that RNAi-dependent epimutation functions as a host-induced regulatory program that remodels fungal gene expression during infection. We identify a set of brain-induced epimutation loci and demonstrate that deletion of one of these genes, the MAPK-like kinase *bepA,* enhances migration across a human BBB model. These findings suggest that RNAi-mediated epimutation is not simply a mechanism of phenotypic diversification under stress, but a dynamic strategy through which host microenvironments can reversibly reprogram fungal regulatory networks to facilitate tissue invasion.

This *in vivo* RNAi-based epimutation is also organ-associated. Previous work showed that RNAi-dependent drug-resistance epimutations can be induced and reverted during murine infection, with pronounced effects after recovery from the brain compared with other organs (10). We have identified brain-derived fungal cells that accumulated small RNAs at defined *bep* loci, whereas liver-derived and in vitro samples showed minimal signal at these regions. This pattern argues against a generalized activation of RNAi during host passage and instead supports the existence of organ-specific epigenetic programs. We propose that this represents a form of organ-specific epigenetic adaptation, in which distinct host tissues impose selective pressures that are sensed and translated into reversible gene regulatory states.

The CNS presents a highly specialized environment for invading fungi. To reach and colonize this niche, fungal cells must survive in the bloodstream, interact with vascular endothelium, traverse the BBB, and adapt to a tissue environment that differs markedly from peripheral organs in nutrient availability, immune surveillance, oxygen tension, and cellular architecture (17, 22, 23). CNS involvement is also one of the most devastating manifestations of mucormycosis and is associated with poor clinical outcome (24, 25). The enrichment of RNAi-dependent epimutations after brain passage therefore suggests that the CNS is a host environment capable of actively shaping fungal regulatory states.

The identification of the *bepA* epimutation provides a mechanistic entry point into this process. BepA encodes a conserved MAPK-like kinase with features characteristic of cell wall integrity MAPKs, including a TEY activation loop. This is significant because MAPK pathways occupy central positions in fungal environmental sensing, coordinating stress responses, morphogenesis, cell wall remodeling, and host interaction (26–28). Reversible silencing of *bepA* could therefore provide an efficient mechanism to remodel multiple downstream processes without requiring stable genetic mutation. In this sense, bepA epimutation highlights the potential for host-associated epimutations to target regulatory nodes that reshape fungal behavior during infection. Our data support a role for BepA in calcineurin-associated stress signaling. Loss of *bepA* altered the response to calcineurin inhibitors, allowing hyphal growth in the presence of FK506 and cyclosporin A. This phenotype is notable because calcineurin controls dimorphic growth and virulence-associated morphogenesis in *Mucorales* (12, 29). In contrast, the *bepA* deletion strain retained normal anaerobic yeast growth, indicating that BepA is not a general regulator of yeast development modulated by PKA (30–32), but rather participates in specific calcineurin-dependent morphogenetic responses. Thus, BepA defines a stress-responsive branch of the calcineurin-associated morphogenetic network that is modulated by RNAi epimutation under CNS-induced stress.

Our analysis links this dimorphic signaling function to a downstream cell wall remodeling program. BepA-dependent genes substantially overlapped with FK506-responsive genes, indicating that BepA controls a subset of calcineurin-associated transcriptional outputs. This genetic response was enriched for functions related to cell wall biosynthesis and remodeling, including chitin synthases and chitin deacetylases. This convergence is highly relevant to mucormycosis pathogenesis. The cell wall is not only a structural scaffold, but also a dynamic interface between fungal cells and the host (33). In the Mucorales, chitin and chitosan are dominant wall components, and changes in their abundance or organization could influence morphology, stress tolerance, immune recognition, and interaction with host barriers. Recent work has shown that dimorphic Mucorales have evolved duplicated gene families whose paralogs are differentially expressed in yeast and mycelial forms, providing a genomic mechanism to modulate fungal physiology to distinct morphotypes (34). In this context, the presence of multiple chitin synthase and chitin deacetylase genes among the BepA-dependent transcriptional outputs raises the possibility that BepA regulates part of this morphotype-associated cell wall repertoire. We therefore propose that *bepA* silencing shifts cell wall regulatory programs in a manner that favors host tissue interaction and penetration.

The BBB model provides functional support for this model and represents an important methodological advance. To our knowledge, this study establishes the first human BBB transwell platform optimized to quantify transendothelial migration of a filamentous fungus. Previous BBB systems have provided powerful insight into CNS invasion by yeast pathogens such as *C. neoformans* and *Candida albicans* (17, 18, 35, 36), but comparable experimental platforms for filamentous fungi have remained limited. This gap is particularly important for mucormycosis, where angioinvasion, endothelial interaction, and CNS dissemination are defining features of severe disease. With this platform, we found that loss of *bepA* enhanced migration across human brain endothelial monolayers without causing major disruption of TEER values. This result is important because it suggests that increased BBB crossing is not simply a consequence of nonspecific barrier destruction. Instead, *bepA* silencing may enhance fungal-endothelial interaction and provide a functional explanation for the enrichment of *bepA*-associated small RNAs in brain-derived fungal populations. Brain-induced silencing of *bepA* may therefore generate a regulatory state that improves the ability of fungal cells to traverse the barrier required for CNS invasion.

Although loss of *bepA* increased migration across the *in vitro* BBB model and was associated with higher brain fungal burdens, neither deletion nor overexpression significantly altered host survival under the conditions examined. This apparent dissociation could reflect distinct roles of BepA during BBB traversal and subsequent survival or persistence within the brain parenchyma. One possibility is that loss of BepA facilitates barrier crossing while imposing costs on other infection-related functions, potentially offsetting its contribution to overall virulence. However, such a trade-off remains hypothetical, as impaired survival within brain tissue has not been demonstrated. Whole-brain fungal burdens integrate tissue entry, subsequent fungal growth, and host clearance and therefore cannot distinguish these contributions. Furthermore, fungal burden and host survival were assessed in different mouse backgrounds, limiting direct comparisons between these outcomes. Thus, the absence of a detectable survival difference does not exclude a role for BepA in specific stages of infection, but the relationship between its effects on barrier traversal, intracerebral persistence, and disease severity remains unresolved.

More broadly, our findings suggest that RNAi-dependent epimutation provides filamentous fungal pathogens with a reversible route to adaptive phenotypic variation during host colonization. Unlike stable genetic mutation, epimutation enables rapid, locus-specific changes in gene expression while preserving the capacity to revert when selective pressures change. This concept may extend beyond *Mucorales*. Although the molecular mechanisms differ among fungal pathogens, several systems illustrate how reversible, non-genetic regulatory states can shape infection. In *C. albicans*, epigenetic white-opaque and white-gray-opaque switching generates heritable cell states that differ in metabolism, immune interaction, tissue preference, and virulence (37, 38). In *Nakaseomyces glabratus* (formerly *Candida glabrata*), subtelomeric silencing regulates adhesin families that control adherence to host cells (39). In *C. neoformans*, host-induced developmental transitions such as titan cell formation remodel cell size, capsule, cell wall architecture, stress resistance, and interaction with phagocytes during infection (40, 41). Our study places RNAi-dependent epimutation within this broader framework of fungal regulatory plasticity, while revealing a distinct mechanism in which a host organ induces small RNA-mediated silencing of endogenous genes linked to invasion. Thus, brain-induced *bepA* epimutation supports a broader principle: invasive fungi can exploit reversible regulatory states to adapt to local host microenvironments, and these states may actively influence tissue tropism, barrier crossing, and disease progression. We propose that organ-specific epigenetic adaptation represents a key regulatory mechanism of fungal pathogenesis, in which host tissues actively shape fungal phenotypes through reversible RNAi-mediated gene silencing. Understanding how these epigenetic states arise, persist, and revert may reveal new vulnerabilities in mucormycosis, particularly during CNS invasion.

## Materials and methods

### Fungal strains and culture conditions

The *M. circinelloides* mutant strains used in this study were derived from MIN6, an auxotrophic derivative of *M. circinelloides 1006PhL* (42). All strains used in this study are listed in Table S1. Strains were routinely cultured on yeast extract-peptone-dextrose (YPD) medium at 26°C unless otherwise indicated. For sporulation, cultures were grown on YPD agar plates, and sporangiospores were harvested in sterile distilled water and quantified using a hemocytometer. Media supplemented with antifungal drugs or signaling inhibitors were prepared immediately before use. For anaerobiosis growth, strains were grown in liquid YPD for 12h under anaerobic conditions using an anaerobic jar with Anaerobe Container System Sachets (Becton Dickinson Ireland).

### Generation of *bepA* mutant and overexpression strains

Deletion and overexpression strains were generated using parental strain MIN6 by homologous recombination following protoplast generation and electroporation as previously described (42). Gene replacement cassettes containing approximately 1 kb flanking regions were assembled by overlap PCR, the endogenous *bepA* locus was replaced with the *pyrF* marker, and transformants were selected on selective medium without uridine. Correct integration was confirmed by PCR. For *bepA* overexpression, the endogenous *bepA* promoter was replaced with the constitutive *tef1* promoter by homologous recombination. The overexpression cassette was assembled by overlap PCR using homologous regions flanking the targeted promoter region and introduced into MIN6 protoplasts using the same transformation procedure. Transformants were selected for uracil prototrophy, and correct integration of the *tef1* promoter upstream of the *bepA* coding sequence was confirmed by diagnostic PCR. Increased *bepA* transcript abundance in independent overexpression strains was subsequently verified by RT-qPCR. For RT-qPCR analysis, cDNA was synthesized from 1 µg of total RNA using the iScript cDNA Synthesis Kit (Bio-Rad). Quantitative PCR was performed in technical triplicate using a QuantStudio Real-Time PCR System (Applied Biosystems) with SYBR Green PCR Master Mix (Applied Biosystems), according to the manufacturer’s instructions. Relative *bepA* transcript levels were calculated using the ΔΔCT method, with *vma1*, a vacuolar ATPase encoding gene that is constitutively expressed as the endogenous reference. Primer sequences are listed in Table S2.

### RNA purification, library preparation, and sequencing

Fungal colonies recovered from brain and liver of infected mice were plated on YPD agar and grown for 4, 6, or 8 days before RNA purification. Total RNA was extracted from recovered colonies and control plates using the miRNeasy Mini Kit (Qiagen). Small RNA (sRNA) libraries were prepared using the QIAseq miRNA Library Kit (Qiagen) and sequenced on an Illumina NextSeq 500 High-Output system to generate 75-bp single-end reads.

For transcriptome sequencing, wild-type (WT), *bepA* deletion (*bepA*Δ), and *bepA* overexpression (*bepA*OE) strains were grown for 16 h in liquid YPD in the absence or presence of FK506 (1 mg/L). Total RNA was purified using the RNeasy Mini Kit (Qiagen). rRNA-depleted libraries were prepared using the Illumina Stranded Total RNA Prep with Ribo-Zero Gold and *Mucor circinelloides* rDNA-specific probes previously designed (7) and sequenced on an Illumina NovaSeq 6000 platform to generate 150-bp paired-end reads.

### sRNA data analysis

Raw sRNA sequencing reads were quality assessed using FastQC v0.11.7 before and after preprocessing. Adapter and low-quality sequences were removed using Trim Galore! v0.6.2. Reads with a Phred quality score ≤20 or a total length ≤13 nt were discarded, and 3′ adapter sequences were removed when an overlap of at least 2 nt was detected. Processed reads were aligned to the *M. circinelloides* 1006PhL reference genome and corresponding annotation obtained from FungiDB (dataset DS_8b08c1c31d) using BWA v0.7.8. Gene-level sRNA counts were generated from BAM files using HTSeq v0.9.1 and the corresponding FungiDB gene annotation. Reads were counted in a strand-specific manner (--stranded=yes). Reads overlapping annotated exons were assigned to genes using the gene_id attribute in union mode, while multimapping reads were excluded. Individual count files were combined to generate a gene-by-sample raw count matrix for downstream differential sRNA abundance analysis.

Differential sRNA accumulation was analyzed using the limma-voom framework (limma v3.38.3). Low-abundance loci were filtered before analysis, library composition was normalized using the trimmed mean of M-values (TMM) method, and counts were transformed to log2 counts per million (log2CPM) using voom. Samples collected at 4, 6, and 8 days were grouped according to experimental condition and treated as replicate observations for the organ-level comparison, yielding three observations each for brain, liver, and control conditions. Linear models were fitted in limma and significance was evaluated using empirical Bayes-moderated statistics. P values were corrected for multiple testing using the Benjamini–Hochberg false discovery rate (FDR) procedure. Loci with FDR ≤0.05 were considered differentially represented; an absolute log2 fold change ≥1 and average abundance ≥1 CPM were used to select strongly differentially represented loci for downstream analyses.

For heatmap visualization, normalized log2CPM values were standardized independently for each locus by subtracting the mean across samples and dividing by the corresponding standard deviation to obtain row-wise Z-scores. Heatmaps were generated in R using pheatmap v1.0.12. Hierarchical clustering of both loci and samples was performed using Manhattan distance and Ward.D agglomerative clustering.

### mRNA data analysis

Newly generated *M. circinelloides* strain 1006PhL raw paired-end mRNA-seq reads were quality assessed using FastQC v0.11.7. Adapter removal and read filtering were performed using fastp v0.23.4 (average Q ≥30 and a minimum read length of 130 nt, with poly-G trimming enabled). Processed reads were aligned to the *M. circinelloides* 1006PhL reference genome and corresponding annotation obtained from FungiDB (dataset DS_8b08c1c31d) using STAR v2.7.11b. Alignments were performed with minimum and maximum intron lengths of 10 and 500 bp, respectively, and coordinate-sorted BAM files were generated using SAMtools v1.10.

Gene-level counts were generated using featureCounts v2.0.6 (Subread package). Paired-end fragments overlapping annotated exons were assigned to genes using the corresponding GTF annotation in reverse-stranded mode. Lowly expressed genes were removed using filterByExpr in edgeR, and library composition was normalized by the TMM method. The normalized count data were transformed using voom, linear models were fitted with limma, and statistical significance was evaluated using empirical Bayes moderation. Pairwise contrasts were generated among the relevant genotype and growth-condition combinations. Normalized voom expression values were used for downstream visualization. Volcano plots and additional transcriptomic visualizations were generated with ggplot2 v3.4.2. For selected genes, expression values from biological replicates were averaged by condition, converted to row-wise Z-scores, and visualized using ComplexHeatmap, with hierarchical clustering of genes while retaining the predefined order of experimental conditions. Gene Ontology enrichment analyses were performed on selected differentially expressed gene sets utilizing FungiDB.

For genomic coverage visualization, CPM-normalized bigWig coverage tracks were generated using bamCoverage from deepTools v3.5.6 with 25-bp bins. Where replicate coverage profiles were combined for visualization, bamCompare v3.5.6 was used to calculate the mean normalized coverage. Tracks were displayed with pyGenomeTracks together with the corresponding genome annotation.

Previously published RNA-seq datasets from *Mucor lusitanicus* strain R7B grown either alone or in coculture with J774A.1 murine macrophages for 5 h (GSE117385) (11) were additionally used to examine transcriptional profiles at selected syntenic loci. Processed reads were aligned to the *M. lusitanicus* MU402 reference genome (Muccir1_3) and corresponding annotation obtained from the JGI MycoCosm portal using STAR, and normalized coverage tracks were generated with bamCoverage for visualization with pyGenomeTracks as described above.

Gene Ontology enrichment analyses were performed on selected differentially expressed gene sets, and heatmaps, volcano plots, and additional transcriptomic visualizations were generated in R.

### Ortholog identification and phylogenetic analyses

BepA-related proteins and fungal MAPKs were identified from proteomes representing the Mucorales *M. circinelloides*, *M. lusitanicus*, *Rhizopus delemar*, *Rhizopus microsporus*, *Lichtheimia corymbifera*, and *Phycomyces blakesleeanus*. To assess the phylogenetic distribution of the identified MAPK lineages outside the Mucorales, the analysis also included *R. irregularis* (Glomeromycota) and *Spizellomyces punctatus* (Chytridiomycota), together with characterized MAPKs from the Dikarya species *C. neoformans*, *Saccharomyces cerevisiae*, *C. albicans*, *Schizosaccharomyces pombe*, *Aspergillus fumigatus*, and *Neurospora crassa*.

The orthogroup containing BepA was identified using OrthoFinder v3.1.5, retaining all paralogous members rather than restricting the analysis to one-to-one orthologs. In parallel, BepA was searched against the combined fungal proteome database using BLASTP v2.17.0+ (E value ≤1 × 10^−5^), and profile hidden Markov models representing the major fungal MAPK families (Mpk1/Slt2 cell wall integrity, Hog1 stress response, and Fus3/Kss1 pheromone/filamentation pathways) were generated from curated reference proteins and searched against all proteomes using HMMER v3.4. Candidate proteins were retained based on membership in the BepA orthogroup, significant similarity to the MAPK profiles, or BepA similarity within Mucorales. For profile-based selection, domain hits with an independent E value ≤1 × 10^−20^ and ≥65% profile coverage were retained using relative profile-score thresholds of 0.70 for the general MAPK profile and 0.75 for family-specific profiles. Additional BepA-like proteins in Mucorales were retained when BLASTP hits had an E value ≤1 × 10^−20^, ≥60% query coverage, and a bitscore ≥60% of the best BepA hit within the corresponding species. Protein kinase domains were identified using HMMER v3.4 and the Pfam Pkinase profile (PF00069) from the Pfam-A database. The highest-scoring kinase domain was extracted from each candidate protein, retaining domains of at least 120 amino acids with an independent E value ≤1 × 10^−5^. Curated experimentally characterized fungal MAPKs were included as phylogenetic references for the Mpk1/Slt2, Hog1, and Fus3/Kss1 families. Exact full-length sequence duplicates between curated references and proteome-derived candidates were removed, whereas non-identical proteins from the same species were retained to preserve potential paralogs.

Kinase-domain sequences were aligned using MAFFT v7.526 with the L-INS-i strategy (--localpair --maxiterate 1000). The alignment used for the primary phylogenetic analysis was trimmed with ClipKIT v2.12.1 in smart-gap mode. Maximum-likelihood phylogenetic inference was performed with IQ-TREE v3.1.2 using ModelFinder to select the best-fitting amino-acid substitution model, with branch support estimated from 1,000 ultrafast bootstrap and 1,000 SH-aLRT replicates. Robustness of the inferred topology to alignment processing was additionally evaluated using trimAl v1.5.rev1 (automated1) and the untrimmed MAFFT alignment. Trees were treated as unrooted and visualized and annotated with iTOL.

### Drug susceptibility and stress assays

Drug susceptibility was evaluated by spotting serial dilutions of sporangiospores onto YPD agar supplemented with FK506 (1 μg/mL), rapamycin (100 ng/mL), cyclosporin A (CsA, 100 μg/mL), FK506 plus rapamycin (1 μg/mL and 100 ng/mL, respectively), FK506 plus CsA (1 μg/mL and 100 μg/mL, respectively), caspofungin (48 μg/mL), FK506 plus caspofungin (1 μg/mL and 48 μg/mL, respectively), posaconazole (2 or 4 μg/L), or amphotericin B (4 μg/L). Plates were incubated at 26°C and photographed after incubation.

### Western blot analysis

For analysis of MAPK activation, fungal cultures of the wild-type, *bepA*Δ, and *bepA^OE^*strains were grown in YPD and in YPD supplemented with FK506 (1 mg/L) liquid media for 16 hours. Cells were harvested by centrifugation, blotted dry with paper towels, and lyophilized. The dried cells were disrupted by beads beating with silica beads. Protein extraction buffer (5 mM Tris-HCl [pH7.5], 1% sodium deoxycholate, 5 mM sodium pyrophosphate, 100 μM sodium orthovanadate, 50 mM NaF, 0.1% sodium dodecyl sulfate [SDS], 1% Triton X-100, 0.5 mM phenylmethylsulfonyl fluoride [PMSF], and protease inhibitor cocktail tablet [Roche]) was added at a ratio of 1ml per 500 mg of dried cells, followed by an additional round of bead beating. The lysates were centrifuged, and protein concentrations in the resulting supernatants were determined using BCA assay (ThermoFisher). Equal amounts of protein were subjected to 10% SDS-PAGE and Western blot analysis using anti-phospho-p44/42 MAPK (Cell Signaling Technology) and anti-GAPDH (Proteintech) antibody.

### Human blood-brain barrier model

A human blood-brain barrier (BBB) transwell model was established using the human brain microvascular endothelial cell line hCMEC/D3 (Merck), adapting a previously described protocol for *C. neoformans* (17). Briefly, 5 × 10^4^ hCMEC/D3 cells were seeded in EGM-2 medium (Lonza) onto collagen-coated transwell inserts with 8-μm porous membranes (BD Falcon) and maintained at 37°C with 5% CO₂. The following day, the medium was replaced with fresh EGM-2 supplemented with 2.5% human serum (Sigma), and cells were cultured for an additional 4 days. One day before fungal inoculation, the medium was replaced with EGM-2 supplemented with 1.25% human serum. Formation and integrity of the endothelial monolayer were assessed by transendothelial electrical resistance (TEER) using an EVOM2 epithelial voltohmmeter. For BBB crossing assays, 5x10^5^ spores of the wild-type, *bepA*Δ, or *bepA^OE^* strains were inoculated into the upper (blood-side) compartment. Following 24 h incubation at 37°C with 5% CO₂, fungal cells that migrated into the lower (brain-side) compartment were collected and quantified by CFU counting on YPD agar. The migration ratio was calculated by normalizing the recovered CFU of each strain to that of the wild-type strain. TEER was measured immediately before and after fungal inoculation to assess changes in endothelial barrier integrity. PBS-treated inserts were included as uninfected controls.

### Virulence assays, fungal burden, and immune cell profiling

All animal experiments were performed in accordance with protocols approved by the Duke University Institutional Animal Care and Use Committee (IACUC, protocol number A002-21-01). Male mice were used in all experiments. Animals were monitored daily and euthanized according to predefined humane endpoints.

For small RNA sequencing experiments, four-week-old male BALB/c mice (Charles River) weighing 20–25 g were immunosuppressed with cyclophosphamide by intraperitoneal injection (200 mg/kg body weight) 2 days before infection and every 5 days thereafter. Mice were intravenously challenged by retro-orbital injection with 1x10^6^ spores of *M. circinelloides* 1006PhL. Brain and liver were recovered at 48 h post-infection and plated on YPD agar for fungal recovery and subsequent RNA purification.

For survival analyses, groups of five four-week-old male BALB/c mice were immunosuppressed as described above and intravenously challenged by retro-orbital injection with 1x10^6^ spores of 1006PhL WT, two independent *bepA*Δ strains, or two independent *bepA^OE^* strains. Mice were monitored daily for survival.

For fungal burden analyses and immune cell profiling, male C57BL/6J mice (The Jackson Laboratory) were infected intravenously by retro-orbital injection with 1x10^6^ spores of 1006PhL, *bepA*Δ, or *bepA^OE^* strains. Organs were collected at 3 days post-infection. Brain and spleen homogenates were plated on YPD agar for CFU quantification. Immune cell populations were quantified by flow cytometry from the same samples as the fungal burdens. Spleens were mechanically separated through a 40 µm metal mesh, then spun down at 500 x g for 5 min, and resuspended in PBS. Splenocytes were stained with antibodies for 30 minutes with the following antibodies CD45-BUV395 (BD Horizon, Clone 30-F11), SiglecF-BV421 (Biolegend, Clone S17007L), CD11b-Pe-Cy7 (Biolegend, Clone M1/70), Ly-6G-Alexa488 (Biolegend, Clone 1A8), Ly-6C APC-Cy7 (Biolegend, Clone HK1.4). After a final wash and resuspension, counting beads (Precision Count Beads, Biolegend) were added and samples were analyzed on a BD Fortessa-X20 Flow Cytometer equipped with five lasers (UV, 405, 466, 561, and 640 nm excitation). Data analysis was conducted in FlowJo.

## Data Availability

Data availability. sRNA-seq and mRNA-seq data generated in this study will be deposited in GEO and released upon publication. Previously published RNA-seq data from *M. lusitanicus* co-cultured with macrophages are available under GSE117385. The *M. circinelloides* 1006PhL genome and annotation are available at FungiDB (dataset DS_8b08c1c31d).

## Competing interests

The authors declare no competing interest.

## Acknowledgements and funding sources

We thank Anna Floyd Averette for her continuous technical assistance and the Duke University School of Medicine Sequencing and Genomic Technologies Core Facility for sequencing services. We also acknowledge the Duke Compute Cluster for providing computational resources and technical support. This work was supported by the National Institutes of Health (NIH)/National Institute of Allergy and Infectious Diseases (NIAID) grants R01 AI050113-20, AI039115-28, AI170543-05, and AI172541-04. C.P.-A. is supported by the Beatriz Galindo Program (BG24-00103), funded by the Spanish Ministry of Science, Innovation and Universities and co-funded by the University of Murcia. The funders had no role in study design, data collection and analysis, interpretation of results, or the decision to submit the manuscript for publication. J.H. is Co-Director and Fellow of the Canadian Institute for Advanced Research (CIFAR) program Fungal Kingdom: Threats & Opportunities.

## Authors contributions

C.P.A., M.I.N.M., and J.H. designed research; C.P.A., M.I.N.M., E.A.S., Y.C., Z.C., Z.X., and E.C. performed research; Y.S.B. contributed to the design of the blood–brain barrier experiments; Z.X. and E.C. contributed new reagents; C.P.A., M.I.N.M., and E.A.S. analyzed data; C.P.A. and M.I.N.M. prepared figures and wrote the paper; J.H. supervised the research; and all authors reviewed, edited, and approved the final manuscript.

## Notes

### Competing Interest Statement

The authors have declared no competing interest.

## References

1. O. A. Cornely, et al., Global guideline for the diagnosis and management of mucormycosis: an initiative of the European Confederation of Medical Mycology in cooperation with the Mycoses Study Group Education and Research Consortium. Lancet Infect. Dis. 19, e405–e421 (2019).

2. Geneva: World Health Organization, WHO fungal priority pathogens list to guide research, development and public health action, CC BY-NC-S (2022).

3. A. Skiada, et al., Zygomycosis in Europe: Analysis of 230 cases accrued by the registry of the European Confederation of Medical Mycology (ECMM) Working Group on Zygomycosis between 2005 and 2007. Clin. Microbiol. Infect. 17, 1859–1867 (2011).

4. M. M. Roden, et al., Epidemiology and outcome of zygomycosis: A review of 929 reported cases. Clin. Infect. Dis. 41, 634–653 (2005).

5. S. Calo, et al., Antifungal drug resistance evoked via RNAi-dependent epimutations. Nature 513, 555–558 (2014).

6. Z. Chang, R. B. Billmyre, S. C. Lee, J. Heitman, Broad antifungal resistance mediated by RNAi-dependent epimutation in the basal human fungal pathogen *Mucor circinelloides*. PLoS Genet. 15, e1007957 (2019).

7. C. Pérez-Arques, M. I. Navarro-Mendoza, Z. Xu, G. Walther, J. Heitman, RNAi epimutations conferring antifungal drug resistance are inheritable. Nat. Commun. 16, 7293 (2025).

8. M. I. Navarro-Mendoza, C. Pérez-Arques, J. Heitman, Heterochromatin and RNAi act independently to ensure genome stability in Mucorales human fungal pathogens. Proc. Natl. Acad. Sci. U. S. A. 120, e2220475120 (2023).

9. C. Pérez-Arques, et al., A non-canonical RNAi pathway controls virulence and genome stability in Mucorales. PLoS Genet. 16, e1008611 (2020).

10. Z. Chang, J. Heitman, Drug-resistant epimutants exhibit organ-specific stability and induction during murine infections caused by the human fungal pathogen *Mucor circinelloides*. mBio 10, e02579–19 (2019).

11. C. Pérez-Arques, et al., *Mucor circinelloides* thrives inside the phagosome through an Atf-mediated germination pathway. mBio 10, 1–15 (2019).

12. S. C. Lee, A. Li, S. Calo, J. Heitman, Calcineurin plays key roles in the dimorphic transition and virulence of the human pathogenic zygomycete *Mucor circinelloides*. PLoS Pathog 9, e1003625 (2013).

13. J. Ocampo, et al., Protein kinase A regulatory subunit isoforms regulate growth and differentiation in *Mucor circinelloides:* Essential role of *pkaR4*. Eukaryot. Cell 11, 989–1002 (2012).

14. P. R. Kraus, D. S. Fox, G. M. Cox, J. Heitman, The *Cryptococcus neoformans* MAP kinase Mpk1 regulates cell integrity in response to antifungal drugs and loss of calcineurin function. Mol. Microbiol. 48, 1377–87 (2003).

15. Q. Cheng, et al., Molecular architecture of chitin and chitosan-dominated cell walls in zygomycetous fungal pathogens by solid-state NMR. Nat. Commun. 15, 8295 (2024).

16. M. D. Lenardon, C. A. Munro, N. A. R. Gow, Chitin synthesis and fungal pathogenesis. Curr. Opin. Microbiol. 13, 416–23 (2010).

17. K.-T. Lee, et al., Fungal kinases and transcription factors regulating brain infection in *Cryptococcus neoformans*. Nat. Commun. 11, 1521 (2020).

18. J. Kim, et al., Fungal brain infection modelled in a human-neurovascular-unit-on-a-chip with a functional blood-brain barrier. *Nat*. Biomed. Eng. 5, 830–846 (2021).

19. S. Calo, et al., A non-canonical RNA degradation pathway suppresses RNAi-dependent epimutations in the human fungal pathogen *Mucor circinelloides*. PLoS Genet. 13, e1006686 (2017).

20. F. E. Nicolas, et al., Endogenous short RNAs generated by Dicer 2 and RNA-dependent RNA polymerase 1 regulate mRNAs in the basal fungus Mucor circinelloides. Nucleic Acids Res 38, 5535–5541 (2010).

21. F. E. Nicolás, et al., The RNAi machinery controls distinct responses to environmental signals in the basal fungus *Mucor circinelloides*. BMC Genomics 16, 237 (2015).

22. J. V Forrester, P. G. McMenamin, S. J. Dando, CNS infection and immune privilege. Nat. Rev. Neurosci. 19, 655–671 (2018).

23. Y. Wu, et al., Microglia and amyloid precursor protein coordinate control of transient *Candida cerebritis* with memory deficits. Nat. Commun. 10, 58 (2019).

24. D. P. Kontoyiannis, T. J. Walsh, Mucormycosis. N. Engl. J. Med. 394, 684–698 (2026).

25. M. Hoenigl, et al., The emergence of COVID-19 associated mucormycosis: a review of cases from 18 countries. The Lancet Microbe 3, e543–e552 (2022).

26. J. R. Rapala, F. A. Davis, A. Larkin, T. R. O’Meara, Pathogenic fungal sensing and responses to stressful host environments. Annu. Rev. Microbiol. 79, 449–473 (2025).

27. E. Román, D. M. Arana, C. Nombela, R. Alonso-Monge, J. Pla, MAP kinase pathways as regulators of fungal virulence. Trends Microbiol. 15, 181–90 (2007).

28. K. Dichtl, S. Samantaray, J. Wagener, Cell wall integrity signalling in human pathogenic fungi. Cell. Microbiol. 18, 1228–38 (2016).

29. S. C. Lee, et al., Calcineurin orchestrates dimorphic transitions, antifungal drug responses and host-pathogen interactions of the pathogenic mucoralean fungus *Mucor circinelloides*. Mol. Microbiol. 97, 844–865 (2015).

30. S. Vellanki, et al., A novel resistance pathway for calcineurin inhibitors in the human-pathogenic mucorales *Mucor circinelloides*. mBio 11, e02949–19 (2020).

31. A. M. Wolff, K. F. Appel, J. B. Petersen, U. Poulsen, J. Arnau, Identification and analysis of genes involved in the control of dimorphism in *Mucor circinelloides* (syn. *racemosus*). FEMS Yeast Res. 2, 203–213 (2002).

32. J. Ocampo, et al., A subunit of protein kinase a regulates growth and differentiation in the fungus *Mucor circinelloides*. Eukaryot. Cell 8, 933–944 (2009).

33. N. A. R. Gow, M. D. Lenardon, Architecture of the dynamic fungal cell wall. Nat. Rev. Microbiol. 21, 248–259 (2023).

34. G. Tahiri, et al., Coordinated gene family evolution shapes the genome of dimorphic Mucorales. Nat. Commun. 17 (2026).

35. K. Vu, B. Weksler, I. Romero, P.-O. Couraud, A. Gelli, Immortalized human brain endothelial cell line HCMEC/D3 as a model of the blood-brain barrier facilitates in vitro studies of central nervous system infection by *Cryptococcus neoformans*. Eukaryot. Cell 8, 1803–7 (2009).

36. A. Y. Jong, M. F. Stins, S. H. Huang, S. H. Chen, K. S. Kim, Traversal of *Candida albicans* across human blood-brain barrier in vitro. Infect. Immun. 69, 4536–44 (2001).

37. R. E. Zordan, D. J. Galgoczy, A. D. Johnson, Epigenetic properties of white-opaque switching in *Candida albicans* are based on a self-sustaining transcriptional feedback loop. Proc. Natl. Acad. Sci. U. S. A. 103, 12807–12 (2006).

38. L. Tao, et al., Discovery of a “white-gray-opaque” tristable phenotypic switching system in *Candida albicans*: roles of non-genetic diversity in host adaptation. PLoS Biol. 12, e1001830 (2014).

39. A. De Las Peñas, et al., Virulence-related surface glycoproteins in the yeast pathogen *Candida glabrata* are encoded in subtelomeric clusters and subject to RAP1- and SIR-dependent transcriptional silencing. Genes Dev. 17, 2245–58 (2003).

40. J. N. Crabtree, et al., Titan cell production enhances the virulence of *Cryptococcus neoformans*. Infect. Immun. 80, 3776–85 (2012).

41. C. Cao, et al., Ubiquitin proteolysis of a CDK-related kinase regulates titan cell formation and virulence in the fungal pathogen *Cryptococcus neoformans*. Nat. Commun. 13, 6397 (2022).

42. M. I. Navarro-Mendoza, et al., Alternative ergosterol biosynthetic pathways confer antifungal drug resistance in the human pathogens within the *Mucor* species complex. mBio 15, e0166124 (2024).

