## Supplementary Information for "Host brain infection induces RNAi epimutation in *Mucor circinelloides* and enhances blood-brain barrier invasion"

[1]Department of Molecular Genetics and Microbiology, Duke University School of Medicine, Durham, NC 27710, USA. [2]Department of Genetics and Microbiology, Faculty of Biology, University of Murcia, Murcia, 30100, Spain. [3]Department of Biotechnology, College of Life Science and Biotechnology, Yonsei University, Seoul, Republic of Korea. [4]Department of Integrative Immunobiology, Duke University School of Medicine, Durham, NC 27710, USA.

[#]Equally contributing authors. *To whom correspondence may be addressed

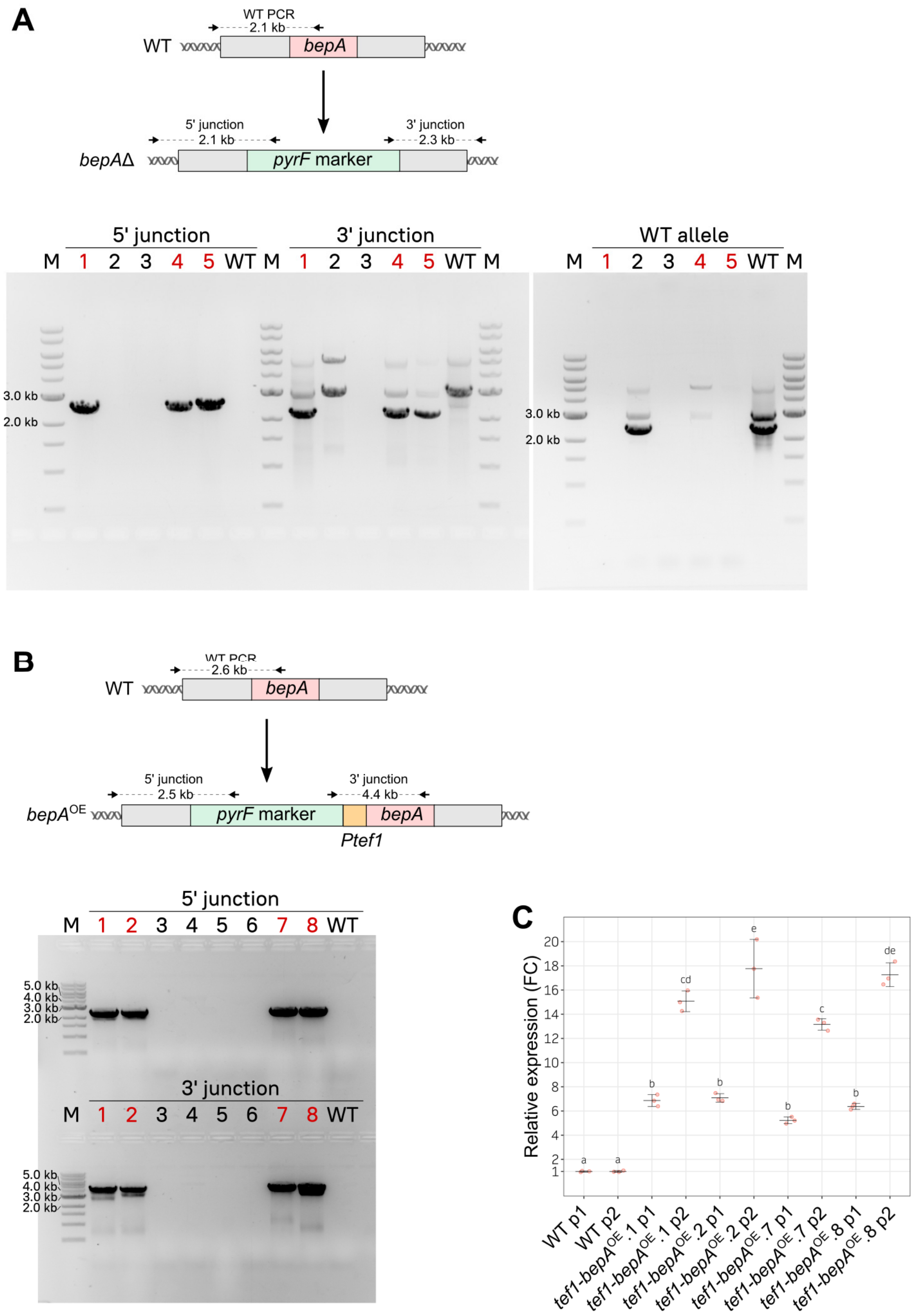


**Figure S1. Generation and validation of *bepA* deletion and overexpression strains in *M. circinelloides*. (A)** Diagram (top) and agarose gel images (bottom) showing replacement of the *bepA* coding region with the *pyrF* selectable marker to generate *bepA*Δ mutants. Arrows indicate the annealing regions of the primers used to confirm correct integration by two PCR reactions corresponding to the 5′ and 3′ junctions. Mutant homokaryosis was confirmed by absence of the wild-type (WT) allele. Lanes are numbered to indicate transformant isolates, M denotes the DNA molecular size marker, and homokaryotic mutants generated by independent transformation experiments and selected for further analysis are highlighted in red. **(B)** Diagram (top) and agarose gel images (bottom) showing generation of *bepA* overexpression strains by integration of the constitutive *tef1* promoter (*Ptef1*) upstream of *bepA*. Correct integration was confirmed by PCR amplification of the 5′ and 3′ junctions. Transformants selected for further analysis are highlighted in red. **(C)** Relative *bepA* transcript levels in independent *Ptef1-bepA*$^{OE}$ strains determined by RT-qPCR using two independent primer pairs (p1 and p2). Fold-change expression was calculated relative to the WT strain and normalized to *vma1*, a vacuolar ATPase encoding gene that is constitutively expressed. Individual points represent the three technical replicates for each strain and primer pair. Significantly distinct values are indicated with different letters and were calculated by one-way ANOVA and Tukey's tests.

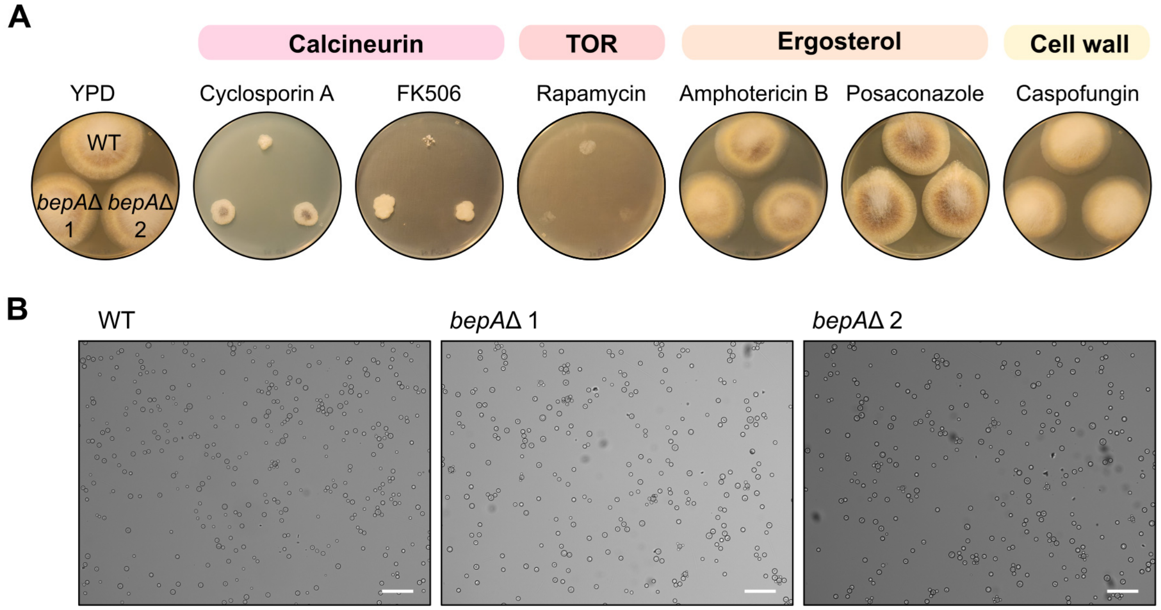


**Figure S2. Growth phenotypes of *bepA* deletion mutants under antifungal stress and anaerobic conditions. (A)** Growth of wild-type (WT) and two independent *bepA* deletion mutants (*bepA*Δ1 and *bepA*Δ2) on YPD control medium and in the presence of cyclosporin A, FK506, rapamycin, amphotericin B, posaconazole, or caspofungin. Compounds are grouped according to their primary cellular targets or pathways. **(B)** Representative light microscopy images of WT, *bepA*Δ1, and *bepA*Δ2 cells after 12 h of growth in liquid YPD under anaerobic conditions. Scale bars, 50 μm.

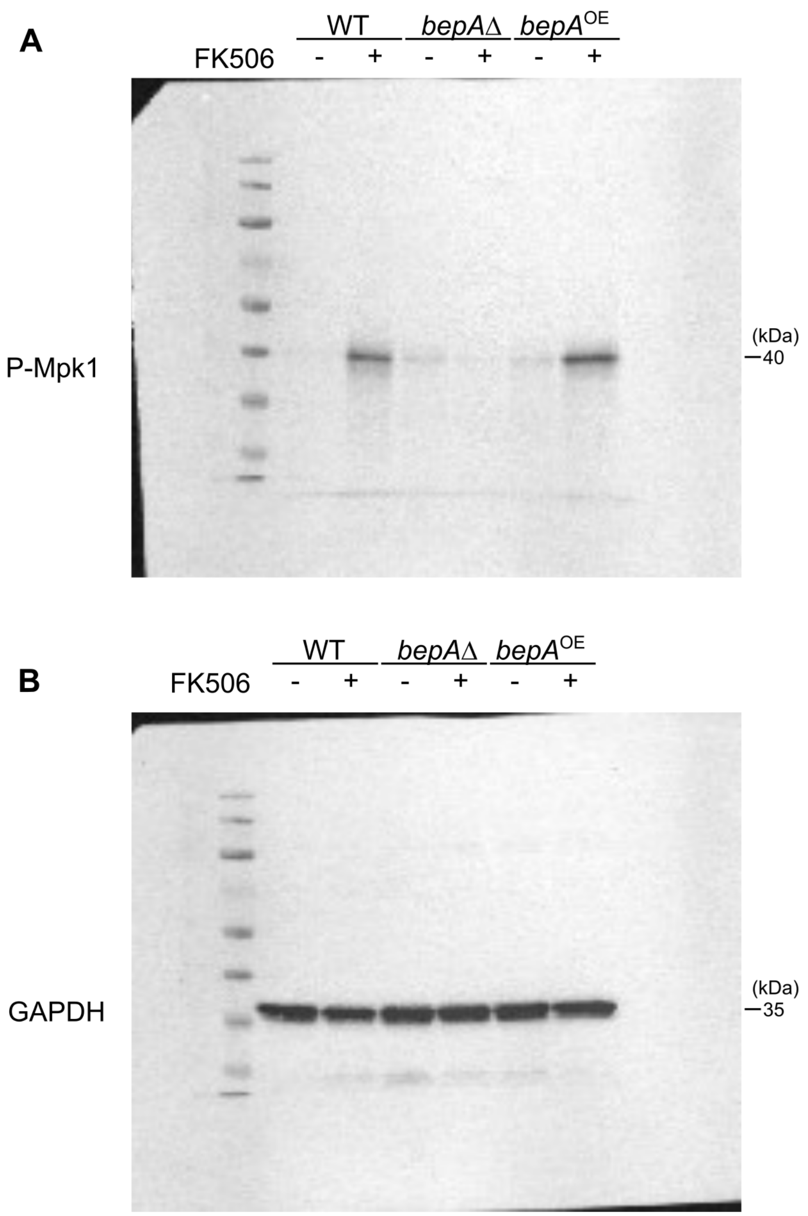


**Figure S3. Uncropped Western blot membranes for analysis of Mpk1 phosphorylation.**

Uncropped Western blot membrane images corresponding to the experiment shown in Fig. 2B. Wild-type (WT), *bepA*Δ, and *bepA*$^{OE}$ strains were grown in the absence (−) or presence (+) of FK506. **(A)** Membrane probed with an anti-phospho-p44/42 MAPK antibody to detect phosphorylated Mpk1 (labeled P-Mpk1 in the panel), with the band used for analysis indicated at approximately 40 kDa. **(B)** Corresponding membrane probed for GAPDH, used as the loading control, with the band detected at approximately 35 kDa. Molecular weight markers are shown on the left of each membrane.

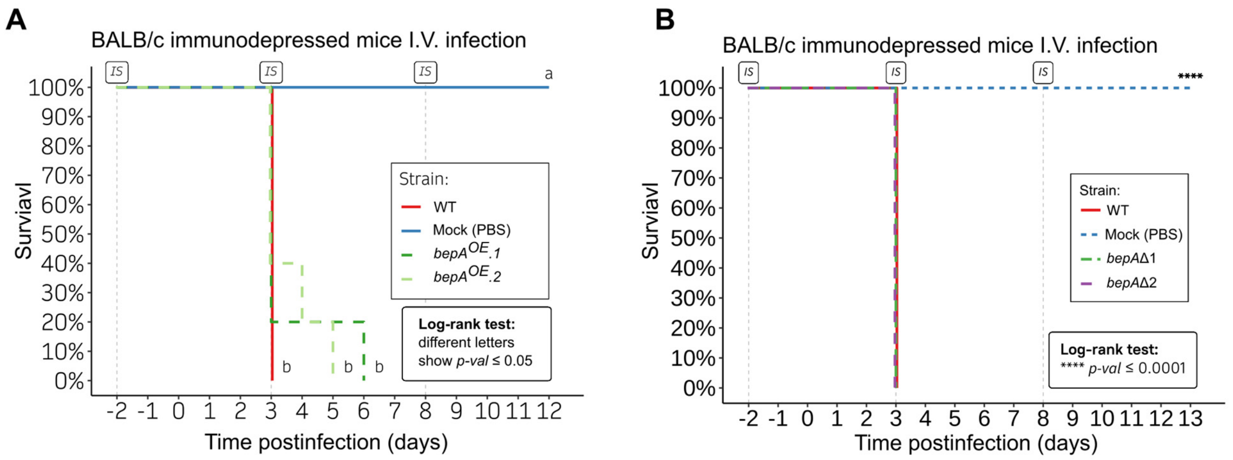


**Figure S4. Survival of BALB/c mice infected with *bepA* mutant strains. (A)** Survival of immunodepressed BALB/c mice following intravenous infection with WT and two independent *bepA*$^{OE}$ strains (*bepA*$^{\mathrm{OE}}$.1 and *bepA*$^{\mathrm{OE}}$.2). Mock-infected animals (PBS) are included as controls. *IS* indicates immunosuppression treatment. Survival differences were assessed by log-rank test, with different letters indicating $p \leq 0.05$. **(B)** Survival of immunodepressed BALB/c mice following intravenous infection with WT and two independent bepA deletion mutants (*bepA*Δ.1 and *bepA*Δ.2). Mock-infected animals are shown as controls. Statistical significance was determined by log-rank test (**** $p \leq 0.0001$).

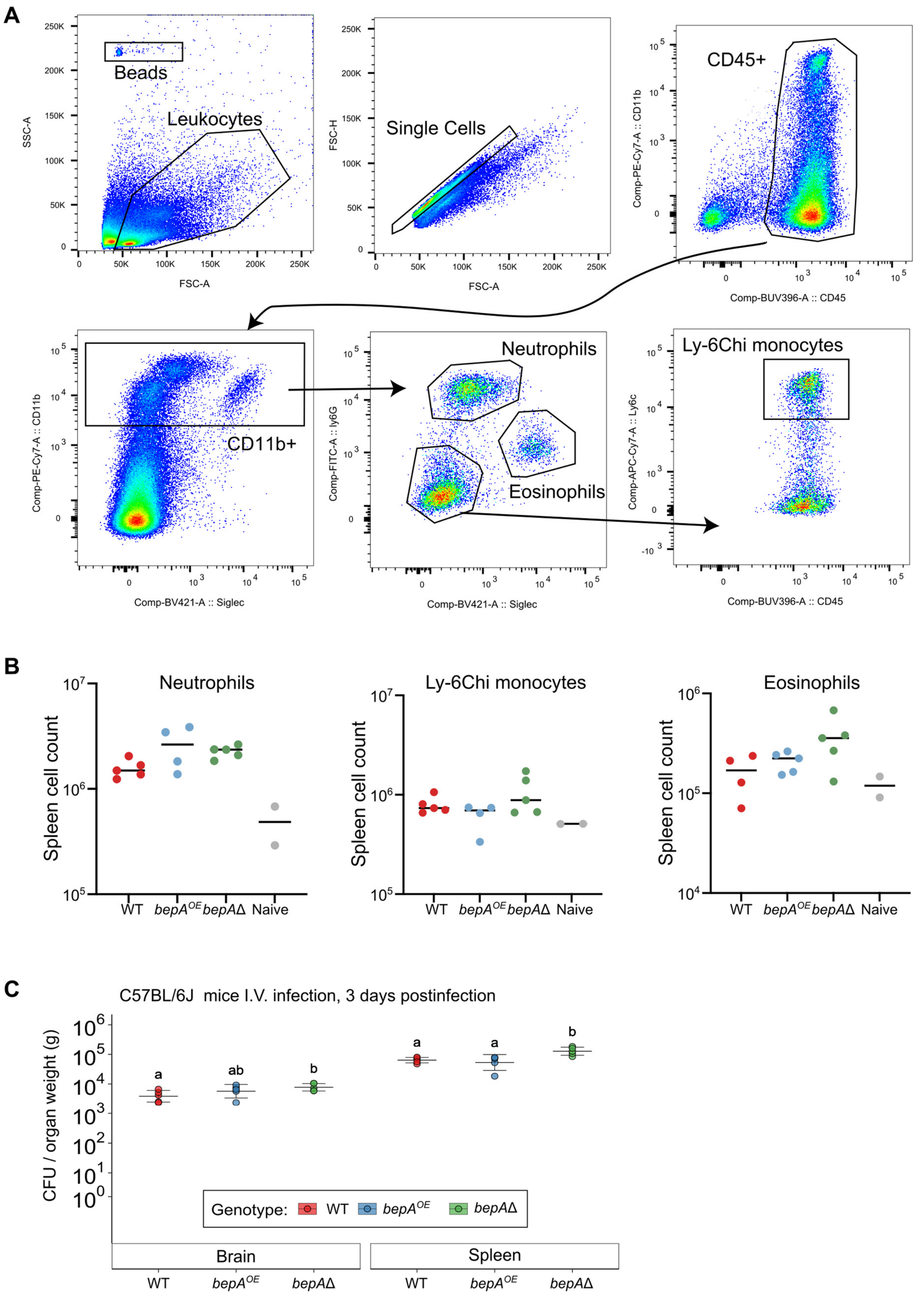
A
Beads
Leukocytes
SSC-A
FSC-A
Single Cells
FSC-H
CD45+
Comp-PE-Cy7-A :: CD11b
Comp-BUV396-A :: CD45
CD11b+
Comp-BV421-A :: Siglec
Neutrophils
Eosinophils
Comp-FITC-A :: ly6G
Ly-6Chi monocytes
Comp-APC-Cy7-A :: Ly6c
B
Neutrophils
Ly-6Chi monocytes
Eosinophils
Spleen cell count
WT
bepA^OE
bepAΔ
Naive
C
C57BL/6J mice I.V. infection, 3 days postinfection
CFU / organ weight (g)
Genotype: WT bepA^OE bepAΔ
Brain
Spleen

64

**Figure S5. Immune cell profiling and fungal burden of *bepA* mutant strains in C57BL/6J mice.** **(A)** Representative flow cytometry gating strategy used to identify splenic myeloid cell populations. Leukocytes were identified based on forward- and side-scatter properties, followed by singlet gating and selection of CD45+ cells. CD11b+ cells were subsequently analyzed to identify neutrophils, eosinophils, and Ly-6Chi inflammatory monocytes based on Ly-6G, Siglec-F, and Ly-6C expression. Counting beads were used to determine absolute cell numbers. **(B)** Absolute numbers of neutrophils, Ly-6Chi inflammatory monocytes, and eosinophils in spleens from mice infected intravenously with wild-type (WT), $bepA^{OE}$ overexpression, or *bepA*Δ deletion strains. Naïve, PBS-injected mice are shown as controls. Each dot represents one animal, and horizontal bars indicate the mean. **(C)** Fungal burden in brain and spleen of C57BL/6J mice at 3 days post-infection following intravenous inoculation with WT, $bepA^{OE}$, or *bepA*Δ strains. Fungal burden is expressed as colony-forming units (CFU) per gram of organ. Each dot represents one mouse, and bars indicate mean ± SD. Different letters indicate statistically significant differences between groups, whereas groups sharing at least one letter are not significantly different ($P < 0.05$; one-way ANOVA).

**Table S1. Strains used in this study**

| Strain | Species | Genotype | Reference |
|---|---|---|---|
| 1006PhL | *M. circinelloides* | wildtype | (42) |
| MIN6 | *M. circinelloides* | *pyrF*-59Δ | (42) |
| MIN19 | *M. circinelloides* | Independent *bepA* deletion mutant; *bepA* replaced with *pyrF* | This study |
| MIN20 | *M. circinelloides* | Independent *bepA* deletion mutant; *bepA* replaced with *pyrF* | This study |
| MNM58 | *M. circinelloides* | Independent *bepA* overexpression strain; constitutive *tef1* promoter | This study |
| MNM60 | *M. circinelloides* | Independent *bepA* overexpression strain; constitutive *tef1* promoter | This study |

**Table S2. Primers used in this study[1]**

| Name | Sequence | Description |
|---|---|---|
| JOHE51498 | AGAATGCCAGACCTGAATTTTTGG | *pyrF* PCR |
| JOHE51499 | AATAGTAATACCCTCTGCCAACGG | *pyrF* PCR |
| JOHE52594 | CGCCCATGACCTATCAGGAC | *bepA* overexpression. WT allele PCR |
| JOHE52595 | ccaaaaattcaggtctggcattctCAAGCCACGGTTCATGGTTG | *bepA* overexpression, upstream flank amplification; overlap with *pyrF* |
| JOHE52596 | ccgttggcagagggtattactattTGTGGGTCTCGGATCGTATG | *tef1* promoter, overlap with *pyrF* |
| JOHE52597 | gtgacataatttgtgtgtgtcatTGCTATGTAAGAAAGATCTGTTAG | *tef1* promoter, overlap with *bepA* |
| JOHE52598 | ATGACACACACAAATTATGTCAC | *bepA* overexpression, coding-region amplification |
| JOHE52599 | CTGCAGGGGGTATCAATGCC | *bepA* overexpression, coding-region amplification. WT allele PCR |
| JOHE51821 | CGCCCATGACCTATCAGGAC | *bepA* deletion |
| JOHE51822 | ccaaaaattcaggtctggcattctCAAGCCACGGTTCATGGTTG | *bepA* deletion, overlap with *pyrF* |
| JOHE51825 | ccgttggcagagggtattactattGAAGACTGTGCCTGCCAATG | *bepA* deletion, overlap with *pyrF* |
| JOHE51826 | GAGTCCAGCGGTATACGTGAG | *bepA* deletion |
| JOHE51863 | CAGATCGCATGGACTGGACA | *bepA,* 5’ junction PCR |
| JOHE51824 | ACACTAGGCGTTTCAAAGCG | *bepA,* 3’ junction PCR |
| JOHE51503 | GAGCAGCAGCATAGAAAGTACCA | *pyrF,* 5’ junction PCR |
| JOHE51504 | TATCAAGAAGGACTAGGCTTGCC | *pyrF,* 3’ junction PCR |
| JOHE51951 | CCGAAGCCCTTGACACTTTG | *bepA* RT-qPCR, pair 1 |
| JOHE51952 | CTGCAGGGGGTATCAATGCC | *bepA* RT-qPCR, pair 1 |
| JOHE52606 | CTTGATACCGTGCACTCACAAT | *bepA* RT-qPCR, pair 2 |
| JOHE52607 | CTTCAAATCTCGGTGAATGACA | *bepA* RT-qPCR, pair 2 |
| JOHE51665 | AGGGTCAATACACTCTTGAAG | *vma1,* RT-qPCR |
| JOHE51667 | GGAGAACGAACAGGCCAGAG | *vma1,* RT-qPCR |

[1]Every primer shows systematic name, nucleotide sequence (in 5’-3’ direction), and a brief description explaining its use.
